# Artificial Soil (ArtSoil): recreating soil conditions in synthetic plant growth media

**DOI:** 10.1101/2025.10.09.681539

**Authors:** Vera Kaplunova, Houda Alioui, Tobias Griguschies, Paloma Durán, Tobias Lautwein, Sabine Metzger, Eliza Loo

## Abstract

Controlled plant growth in laboratories can be achieved by cultivating plants in sterile or axenic conditions on pre-defined synthetic growth media typically supplemented with sugar. In nature, plants do not receive exogenous sugar supply, form symbiosis with microbes, and plant growth is influenced by soil edaphic factors. Thus, physiological and multi-omic analyses from plants grown on synthetic media will differ from those of soil-grown plants due to the influence of sucrose, and the lack of influence from microbiota and soil edaphic factors on plant growth. The rapid advancements in spatial-omics call for accurate characterization of plants grown in conditions similar to soil. To address the issue, we developed Artificial Soil (ArtSoil), a growth medium containing essential nutrients for plant growth, and aqueous soil extract (ASE) to maintain soil microbiomes and edaphic factors, simultaneously eliminating the need for sugar supplementation in the medium. We compared *Arabidopsis thaliana* grown on conventional media to ArtSoil under various growth conditions, and showed that complex soil microbiota in ArtSoil promote plant growth without physiological side effects induced by sucrose. We demonstrate an application for ArtSoil in single-cell transcriptomics and report microbiota-induced cell-type specificity in immune and nitrogen signaling. We tested ArtSoil with six types of ASEs to present the potential of ArtSoil in decoupling the effects of nutrients from microbiota in plant growth. We conclude that ArtSoil recapitulates the soil environment compared to conventional media, hence, enabling physiologically relevant plant growth.

**Significance statement:** Laboratory plant growth commonly relies on sugar-supplemented media and sterile growth conditions, which introduces physiological artifacts and fails to capture the natural influences of soil microbiota and edaphic factors, leading to analyses that diverge from native conditions. We developed Artificial Soil (ArtSoil), a medium containing aqueous soil extract that supports microbiota and sugar-independent growth, and demonstrate its utility in recapitulating native soil properties and enabling physiologically relevant plant development *in vitro*.

## Introduction

A basic requirement in plant research is a gold standard for plant growth conditions, i.e., reproducible conditions as similar as possible to natural growth. Temperature, humidity, light intensity, and diurnal cycle are usually standardized. Since the soil environment is highly heterogeneous, it is common practice to use synthetic growth media to study the effects of specific parameters on the growth and development of mutants and wild-type plants. Various culture media with defined concentrations of macro- and microelements, pH, and gelling agents have been formulated (Gamborg et al., 1968; Hoagland and Armon, D. I., n.d.; Murashige and Skoog, 1962; Schenk and Hildebrandt, 1972). In most cases, growth media are supplemented with sucrose and are kept sterile. It is intriguing that despite the natural ability of plants to photosynthesize to produce sugars, effective axenic growth requires supplementation of the synthetic media with glucose or sucrose. In natural conditions, plants require their microbiome for normal growth and to thrive under stress (Berendsen et al., 2018; Carrión et al., 2019; Castrillo et al., 2017; Durán et al., 2018; Kwak et al., 2018). By contrast, *in vitro* growth are typically conducted in microbe-free conditions.

Sucrose or glucose supplementation in growth media is essential for efficient *Arabidopsis thaliana* (*A. thaliana*) growth as sucrose stimulates root and shoot growth. Sucrose supplementation is also required for *A. thaliana* seed filling and proper embryo formation (Chen et al., 2015; Miotto et al., 2021; Roycewicz and Malamy, 2012). Despite the wide use of sucrose in plant cultivation media, sucrose toxicity can inhibit radicle and primary root growth (S. Liu et al., 2022) as well as seed germination (Li et al., 2012; Yang et al., 2004). Sucrose supplementation also alters nitrogen and carbohydrate metabolism (Huang et al., 2022; McLaughlin et al., 2023; Tan et al., 2022). Growing *A. thaliana* seedlings on sucrose-supplemented media could mask the growth phenotypes of certain mutants (Chen et al., 2015; Fan et al., 2025). For instance, the sucrose uniporter *sweet11;12* double mutants exhibit root growth retardation when grown on sucrose-free media, a phenotype that is masked or compensated for when the seedlings are grown on sucrose-supplemented media (Chen et al., 2015). Sucrose affects *A. thaliana* throughout the plant’s developmental stages as evidenced by contrasting patterns of defense-related genes induction in 9 vs. 14-day-old seedlings (Siffert and Sasse, 2024). The use of sucrose in growth media, thus, poses a complex uncoupling of authentic plant responses from sucrose-induced responses.

Besides altering root hair growth (Claeijs and Vissenberg, 2022), single-cell analyses revealed that roots of sterile *A. thaliana* seedlings grown on sucrose-supplemented media have higher proportions of developmentally mature cells and altered expression of genes that are highly specific to distinct cell populations (Shulse et al., 2019). Root development studies are limited by ecological accuracy since potential mechanisms of root responses mediated by the soil- and root-associated microbiota are not captured in standard growth media (Dini-Andreote et al., 2025). Due to the importance of microbiota for plant growth and development (Durán et al., 2018; Garrido-Oter et al., 2018), synthetic microbial communities and gnotobiotic plant growth systems have been established (Bodenhausen et al., 2014; Gao et al., 2018; Kremer et al., 2021; Ma et al., 2022). The important discoveries have facilitated the understanding of microbiota function, but it remains crucial to study the role of microbiota and the plant under ecosystem-relevant conditions where the total soil microbial community and soil edaphic factors can be captured (Chesneau et al., 2025). Synthetic growth media affect bacterial fitness for colonizing host plants (Torres et al., 2025). Soil properties including nutrients, pH, and salinity directly affect soil microbial community structure and consequently, the assembly of microbiota (Bakker et al., 2018). Although individual strains and consortia of plant-beneficial microbes have been identified (Daniel et al., 2022; Gouda et al., 2018; Poria et al., 2022; Souza et al., 2015), a major challenge in translating microbiota research to the field is integrating the beneficial strain or consortia with the native microbiota (Liu et al., 2024; Russ et al., 2023; Sessitsch et al., 2019). Gnotobiotic systems often cannot fully capture the diversity and complexity of soil microbial communities, which may lead to overlooking important factors that affect community assembly, e.g., priority effects (Carlström et al., 2019; Fukami, 2015; Wippel et al., 2021) and regionalization of microbiota colonization along the root (Loo et al., 2024).

Extensive efforts taken to inform the scientific community about synthetic community design exemplify the complexity of synthetic community design (Durán et al., 2025; Mehlferber et al., 2024; Northen et al., 2024). One approach to addressing the challenges resulting from using sterile sucrose-supplemented media and the complexity of synthetic communities is to use a sucrose-free medium containing self-equilibrating complex soil microbial communities. Here, we present Artificial Soil (ArtSoil) that enables plant cultivation under soil-nurturing growth conditions while enabling controlled environmental conditions and full access to the roots. ArtSoil is a synthetic growth medium supplemented with an aqueous soil extract (ASE) that includes soil microbiota to support growth, hence eliminating the need for sucrose supplementation in the medium (Figure 1). We optimized and compared plants grown on ArtSoil to synthetic growth media with or without sucrose supplementation under various growth conditions and showed that ArtSoil supported plant growth with higher shoot biomass and complex root architecture without negative side effects introduced by sucrose. We demonstrate an application of ArtSoil in single-cell transcriptomics of *A. thaliana* roots and report cell-type-specificity in root microbiota-induction of defense and nitrogen signalling, as well as possible sucrose-induced artefacts in plant-microbe studies. We challenged ArtSoil with six different ASEs and demonstrated the potential to separate nutrient effects from microbiota promotion of plant growth.

**Figure 1:**
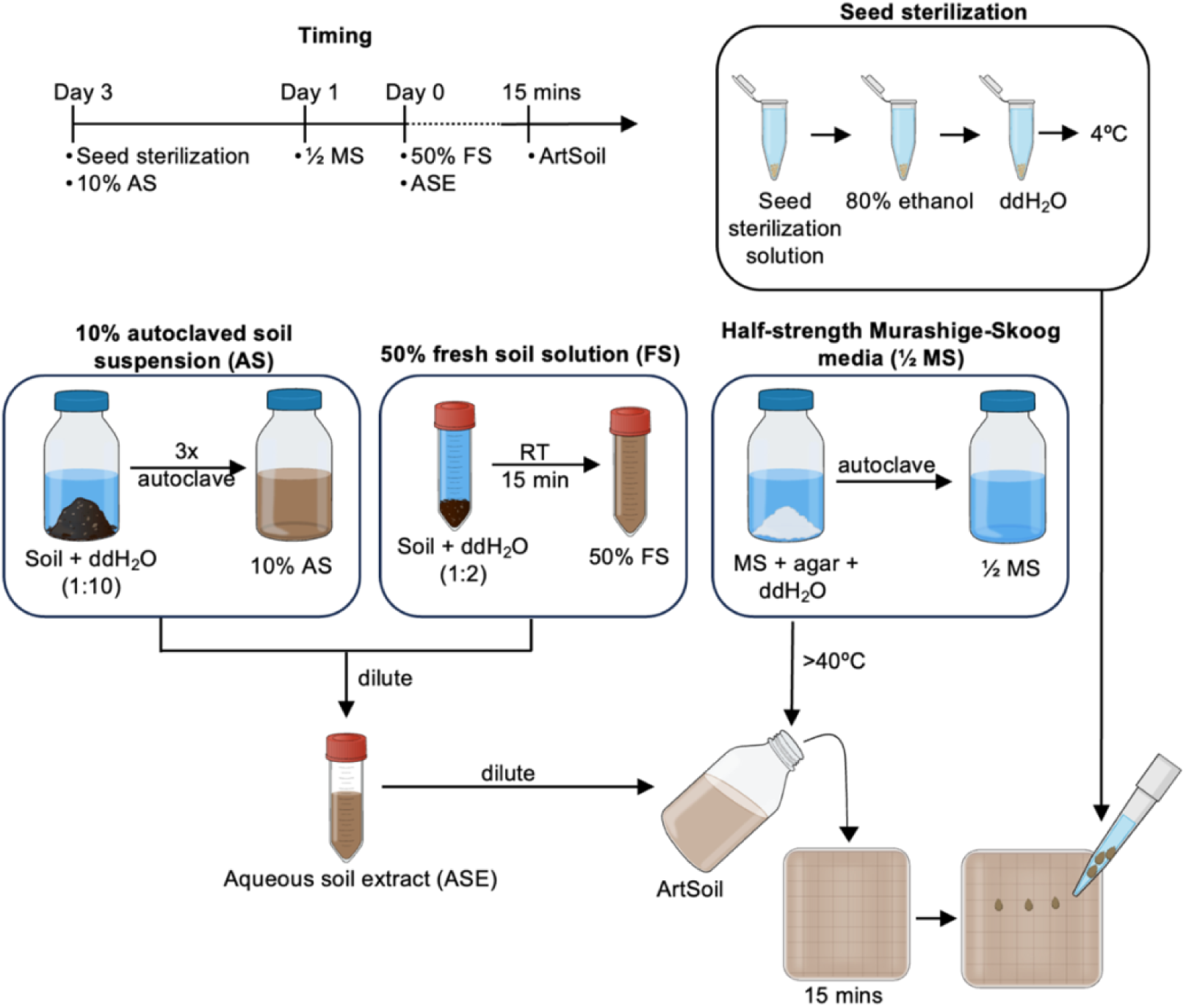
Preparation of ArtSoil. Overview of the steps involved in the preparation of ArtSoil. ArtSoil is composed of three main components, i.e., 10% autoclaved soil solution (10% AS), 50% fresh soil solution (50% FS), and ½ MS without sucrose supplementation. Three days before the assembly of ArtSoil, seeds should be sterilized and 10% AS should be prepared for three consecutive autoclaves. One day before the assembly of ArtSoil plates, ½ MS without sucrose supplementation should be prepared for autoclave. On the day of plate assembly, 50% FS should be prepared and diluted accordingly with 10% AS to produce ASE. The ASE will be added into ½ MS without sucrose supplementation to produce ArtSoil. ArtSoil can be subsequently poured into open or closed Petri dishes and left to cool for the agar to solidify before sowing seeds.

## Results

### Effects of microbial load on root microbiota diversity and plant growth on ArtSoil

The ASE provides soil microbial community to the growth media. Since microbial load and diversity affect the community stability (Cook et al., 2006; Van Elsas et al., 2012; Wippel et al., 2021), it is crucial to determine the optimal dilution of ASE, hence, the amount of microbes used for establishing the microbial community in ArtSoil. The dilution-to-extinction approach was adopted to determine the optimum microbial load for establishing a stable community *in situ* (Cook et al., 2006; Korenblum et al., 2020; Van Elsas et al., 2012). To demonstrate the effects of ASE dilution on plant growth, three dilutions, i.e., 10^-2^, 10^-4^, and 10^-6^ of the Cologne Agricultural Soil (CAS) extract were used for establishing the microbial community in ArtSoil (Method S1). The Shannon-Weaver diversity index was used as a measure of microbial diversity, which typically falls in the range of 1.5 to 3.5 and rarely exceeds 4.5 (Ortiz-Burgos, 2016). CAS had a microbial Shannon-Weaver diversity of 4.3, while roots of CAS-grown *A. thaliana* have a Shannon-Weaver diversity index of 3.4 (Figure 2A). Bulk media from ArtSoil^CAS^ (agar from ArtSoil plates without plants) with 10^-2^, 10^-4^, and 10^-6^ dilutions of CAS had Shannon-Weaver diversity indices of 3.6, 3.0, and 3.0, respectively, indicating reduced microbial diversity in ArtSoil^CAS^ compared to CAS but within the range of expected diversity index. The Shannon-Weaver diversity index of microbes derived from roots of plants grown on ArtSoil^CAS^ inoculated with 10^-2^, 10^-4^, and 10^-6^ of CAS were 2.9, 2.6, and 3.4, respectively. The diversity of microbes from the roots of *A. thaliana* grown on ArtSoil^CAS^ with 10^-6^ dilution of CAS (Shannon-Weaver index 3.4) most closely resembled the diversity of microbes derived from plants grown on soil (Shannon-Weaver index 3.4). The community structures of root-associated microbiota derived from ArtSoil reconstituted the core root microbiota (Bulgarelli et al., 2013; Lundberg et al., 2012) with CAS of 10^-4^ most closely resembling CAS-grown *A. thaliana* (Bulgarelli et al., 2012; Schlaeppi et al., 2014) (Figure 2B).

**Figure 2:**
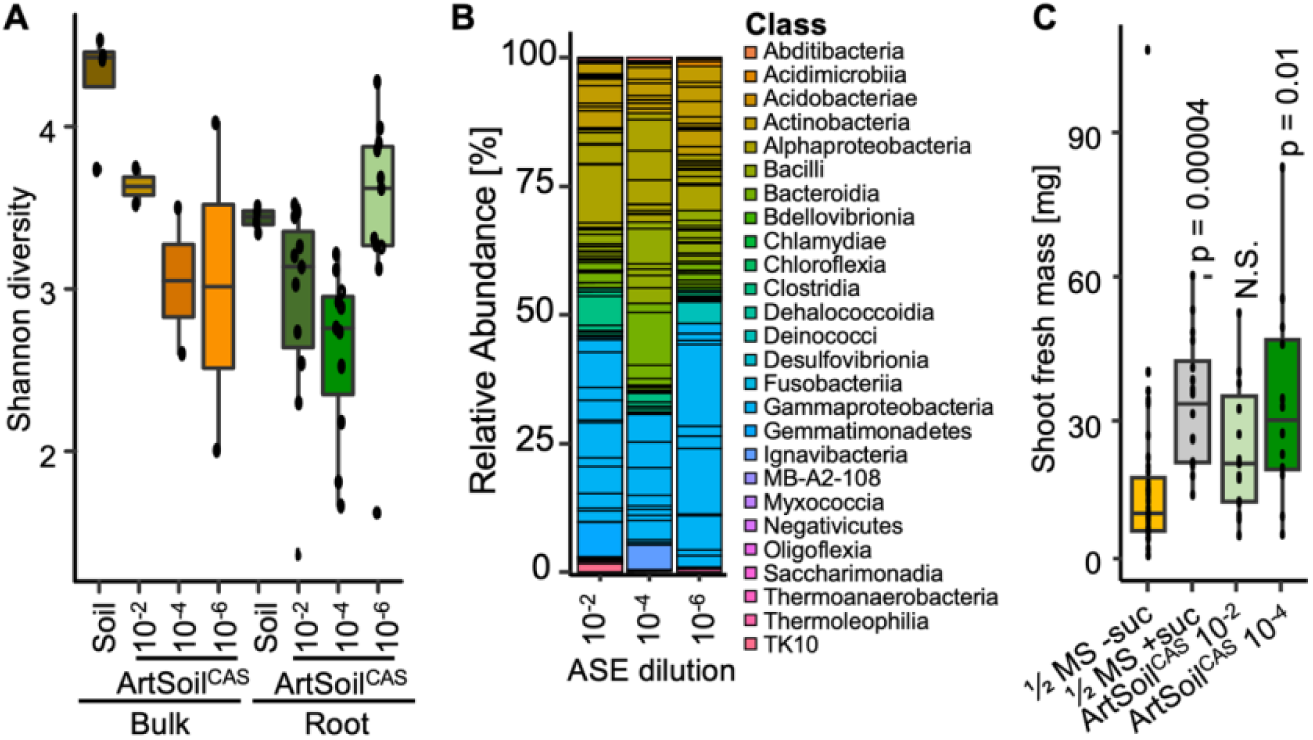
Dilution of ASE influences plant growth on ArtSoil. A) The Shannon-Weaver diversity of root microbiota from 4-week-old *A. thaliana* plants grown on ArtSoil with 10^-2^, 10^-4^, or 10^-6^ dilutions of ASE in a closed system. ‘Bulk’ indicates microbial community derived from soil or ArtSoil media without plants. ‘Root’ indicates microbial community derived from roots of plants grown on soil or ArtSoil with indicated dilutions of ASE. Microbiota profiling performed via quantified via 16S amplicon sequencing. B) Relative abundance of root microbiota derived from 4-week-old *A. thaliana* plants grown on ArtSoil with 10^-2^, 10^-4^, or 10^-6^of ASE. Colours correspond phyla listed in the legend. Each bar represents normalized relative abundance from N > 5. C) Shoot fresh mass of *A. thaliana* seedlings from grown on ArtSoil with 10^-2^ or 10^-4^ compared to sterile ½ MS media. All assays were repeated three times. For Shannon diversity, N > 9; for plant shoot fresh mass, N > 15. P values were calculated via Student’s T-test against ½ MS media without sucrose supplementation (½ MS -suc). N.S.- not significant. The top and bottom boundaries of each boxplot indicate the 1^st^ and 3^rd^ quartiles, the center line indicates the median, and the whiskers represent 1.5× the interquartile range from the 1^st^ and 3^rd^ quartiles.

It should be noted that an imbalance in bacterial abundance may result in plant susceptibility to opportunistic pathogens indicated by reduced plant growth (Ma et al., 2021). Roots of plants from ArtSoil^CAS^ inoculated with 10^-2^ dilution of CAS have a higher Shannon-Weaver diversity index compared to roots from ArtSoil with 10^-4^ dilution of CAS. However, plants grown on ArtSoil^CAS^ inoculated with 10^-2^ dilution of CAS lacked growth promotion conferred by the microbiota compared to sterile ½ MS without sucrose (Figure 2C), indicating possible microbial overload. Since different ASEs have different microbial loads and diversity, users are recommended to test a series of dilutions to determine the optimum dilution of ASE for ArtSoil. Since ArtSoil^CAS^ with a 10^-4^ dilution of ASE was able to reconstitute the root microbiome without plant growth penalty (Figure 2C), subsequent analyses were performed on plants grown on ArtSoil^CAS^ with a 10^-4^ dilution of ASE.

### Influence of ArtSoil on plant development in different growth systems

Plant growth in the closed systems (i.e., sealed Petri dishes) typically causes the accumulation of ethylene and carbon dioxide gases, decreases transpiration due to increased humidity and limited air circulation, reduces cuticle formation, and experiences different utilization of red to far-red light ratio due to wavelengths absorbed by the plastic (Nagel et al., 2020). Consequently, cultivation in a closed system penalizes shoot and root development. ArtSoil alleviates the physiological stresses of plants grown in a closed system, as evidenced by increased shoot biomass and root length (Figure 3A). Petioles of *A. thaliana* grown on half-strength Murashige-Skoog (½ MS) without sucrose supplementation in a closed system were elongated, whereas plants grown on closed system ArtSoil^CAS^ are visibly larger and have extensive root growth and branching (Figure 3A). To test whether ArtSoil could support plant growth beyond physiological stresses introduced in confined growth chambers, ArtSoil was applied to an open plant growth system (Nagel et al., 2020). Compared to the closed system, 4-week-old *A. thaliana* grown on ½ MS without sucrose supplementation on the open system had larger rosette diameters without petiole elongation and a substantially higher root mass (Figure 3B). Plants grown on ArtSoil^CAS^ in an open system had higher shoot biomass and root branching density compared to plants grown on ½ MS without sucrose supplementation (Figure 3B). Thus, ArtSoil^CAS^ promotion of plant growth extends beyond alleviating physiological stresses in closed growth systems.

**Figure 3:**
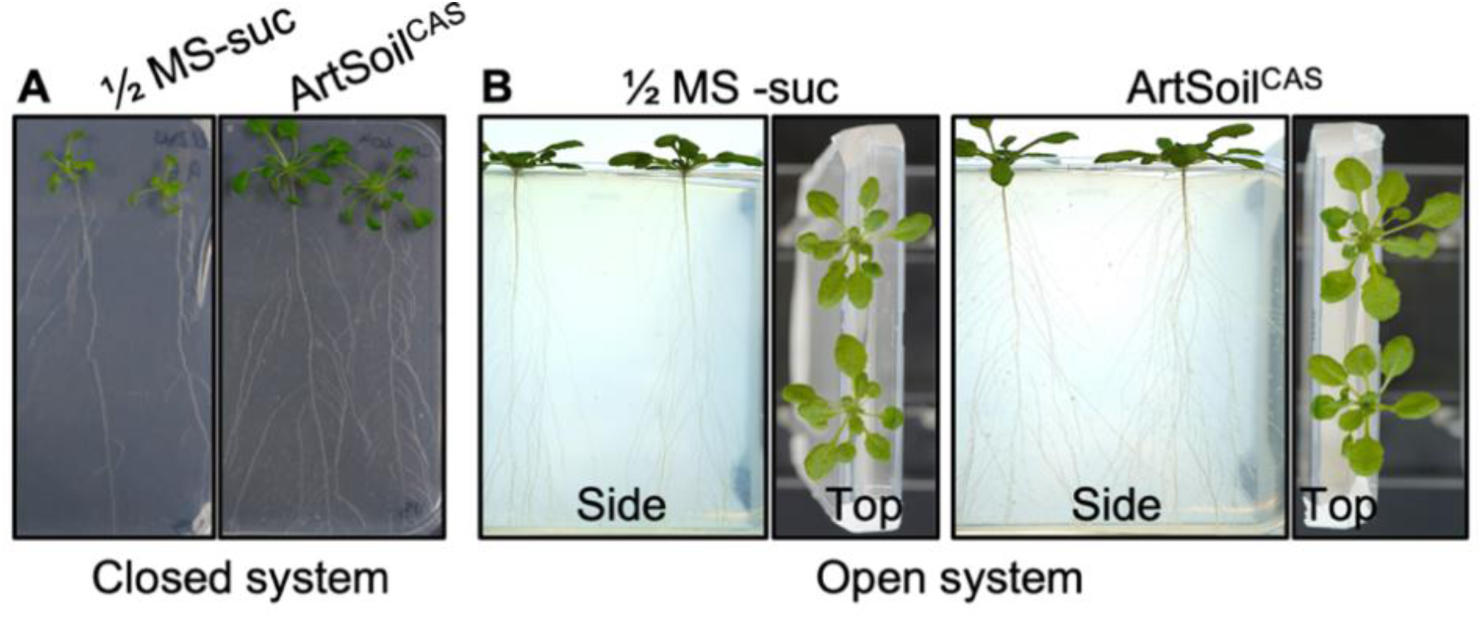
*A. thaliana* growth on ArtSoil in a closed or open system. A) Comparison of phenotypes of 4-week-old *A. thaliana* grown on ½ MS without sucrose supplementation, and ArtSoil with CAS as ASE (ArtSoil^CAS^) in a closed system where shoots and roots were enclosed in sealed Petri dishes. B) Side and top view of 4-week-old *A. thaliana* plants grown on ½ MS without sucrose supplementation, or on ArtSoil^CAS^ in an open system where shoots are exposed to air and roots were enclosed in Petri dishes wrapped with aluminum foil (in darkness).

### ArtSoil does not introduce sucrose-induced artefacts on plant growth

Under sterile growth conditions, sucrose supplementation in growth media promotes *A. thaliana* growth (Kircher and Schopfer, 2023; Pereyra et al., 2022; Roycewicz and Malamy, 2012). Evidently, the shoot fresh mass of 4-week-old *A. thaliana* grown under optimal light intensity (120 μmol m^−2^ s^−1^) on ½ MS media without sucrose supplementation was significantly lower compared to seedlings grown on ½ MS with sucrose supplementation (Figure 4A). The shoot fresh mass of seedlings grown on ArtSoil^CAS^ was significantly higher compared to seedlings grown on ½ MS without sucrose supplementation but not significantly different compared to seedlings grown on ½ MS with sucrose supplementation. Under sub-optimal light intensity, microbiota alleviates plant growth retardation (Hou et al., 2021). Accordingly, the shoot fresh mass of plants grown under low light intensity (25 μmol m^−2^ s^−1^) on ArtSoil^CAS^ was significantly higher than plants grown on ½ MS without sucrose supplementation (Figure 4A). Under low light conditions, exogenous sucrose application could promote root development (Miotto et al., 2021; van Gelderen et al., 2018). The roots of *A. thaliana* seedlings grown at low light intensity (25 μmol m^−2^ s^−1^) on ½ MS without sucrose supplementation or ArtSoil^CAS^ were significantly shorter compared to seedlings grown on ½ MS media with sucrose supplementation (Figure 4B, Figure S1), indicating that ArtSoil^CAS^ does not artificially induce root growth under sub-optimal light conditions. Under optimum light intensity (120 μmol m^−2^ s^−1^), roots of 1- to 3-week-old seedlings grown on ½ MS media without sucrose supplementation or on ArtSoil^CAS^ were significantly shorter compared to roots from seedlings grown on ½ MS media with sucrose supplementation (Figure S1). Notably, roots of 4-week-old seedlings grown on ArtSoil^CAS^ were not significantly different in length compared to roots of seedlings grown on ½ MS media with sucrose supplementation (Figure 4B, Figure S1). Root growth promotion was pronounced after four weeks of growth on ArtSoil^CAS^, a duration that coincides with the duration required for the microbial colonization of the root to stabilize (Edwards et al., 2015; Zhou et al., 2024).

**Figure 4:**
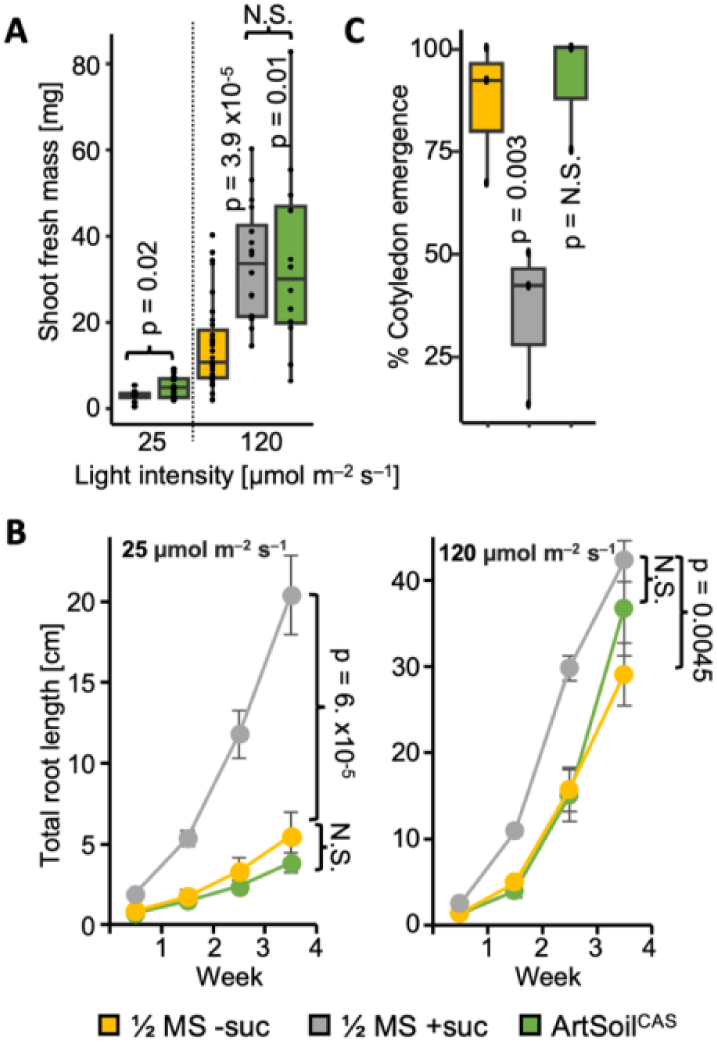
ArtSoil supports *A. thaliana* growth without side effects from sucrose supplementation in synthetic growth media. A) Shoot fresh mass of 4-week-old *A. thaliana* grown on sterile media or in a closed system on ArtSoil^CAS^ under sub-optimal (25 μmol m^−2^ s^−1^) or optimal (120 μmol m^−2^ s^−1^) light conditions. B) Total root length of *A. thaliana* grown on sterile media or in a closed system on ArtSoil^CAS^ for 1 to 4 weeks under sub-optimal (25 μmol m^−2^ s^−1^) or optimal (120 μmol m^−2^ s^−1^) light conditions. Error bars indicate standard errors. C) Percentage cotyledon emergence of *A. thaliana* sown on sterile media or in a closed system on ArtSoil^CAS^ 2 days after sowing. All assays were repeated three times, N > 15. P values were calculated via Student’s T-test against ½ MS without sucrose supplementation (½ MS-suc). N.S.- not significant. The top and bottom boundaries of each boxplot indicate the 1^st^ and 3^rd^ quartiles, the center line indicates the median, and the whiskers represent 1.5× the interquartile range from the 1^st^ and 3^rd^ quartiles.

Although sucrose is beneficial for *in vitro* plant growth, sucrose can also reduce seed germination efficiency (Li et al., 2012). Here, we found that only 35% of *A. thaliana* seeds sown on ½ MS media with sucrose supplementation exhibited cotyledon emergence 2 days after sowing compared to 74% of seeds with cotyledon emergence when sown on ½ MS media without sucrose supplementation (Figure 4C). Similar to ½ MS media without sucrose supplementation, cotyledons emerged from 83% of seeds 2 days after sowing on ArtSoil^CAS^ (Figure 4C). Collectively, our data indicated that ArtSoil enables cultivation of *A. thaliana* without the need for sugar supplementation, thus eliminating possible unphysiological effects of sucrose on plant growth.

### Single-cell analysis of ArtSoil-grown roots reveals cell-type-specific transcriptomic changes induced by microbiota

To investigate whether ArtSoil eliminates sucrose-induced artefacts at the cellular level, single-cell transcriptomics was performed on 7-day-old *A. thaliana* roots grown on ArtSoil. A total of 8000 cells were sequenced. A mean of 23,213 reads per cell, and a median of 2,424 genes per cell were obtained. UMAP clustering and annotation based on previously established marker genes (https://rootcellatlas.org/) revealed 16 transcriptionally distinct clusters representing the major cell types (Figure 5A). The epidermal cell lineage presents six clusters corresponding to trichoblast and atrichoblast cells from all developmental zones. The ground tissue lineage contained three clusters associated with cortex or endodermis cells from the elongation (EZ) and differentiation zone (DZ), forming a continuum that mirrors their radial arrangement *in vivo*. The stele lineage comprised two distinct vascular cell types, including phloem and pericycle clusters in the EZ and DZ, while the root cap lineage included procambium and columella cells. The transitions between clusters indicated that the UMAP captured the gradual changes in gene expression consistent with the developmental and spatial continuity of the root. To quantify the representation of each cell type, the proportion of cell types was calculated (Figure 5B). The epidermal and ground tissue populations were most abundant, followed by the stele and root cap cells. Within the epidermis population, trichoblasts constituted the largest fraction (28.8%), followed by atrichoblasts (16.3%). Cortex and endodermis collectively accounted for most of the ground tissue (17.2%), the stele comprised two main vascular subtypes, pericycle (11.6%) and phloem (5.2%), and procambium and columella cells, comprised approximately 8.3% and 6.4%, respectively. In sum, cell type proportions are representative of root tissues and developmental stages of *A. thaliana* grown under ArtSoil growth conditions.

**Figure 5:**
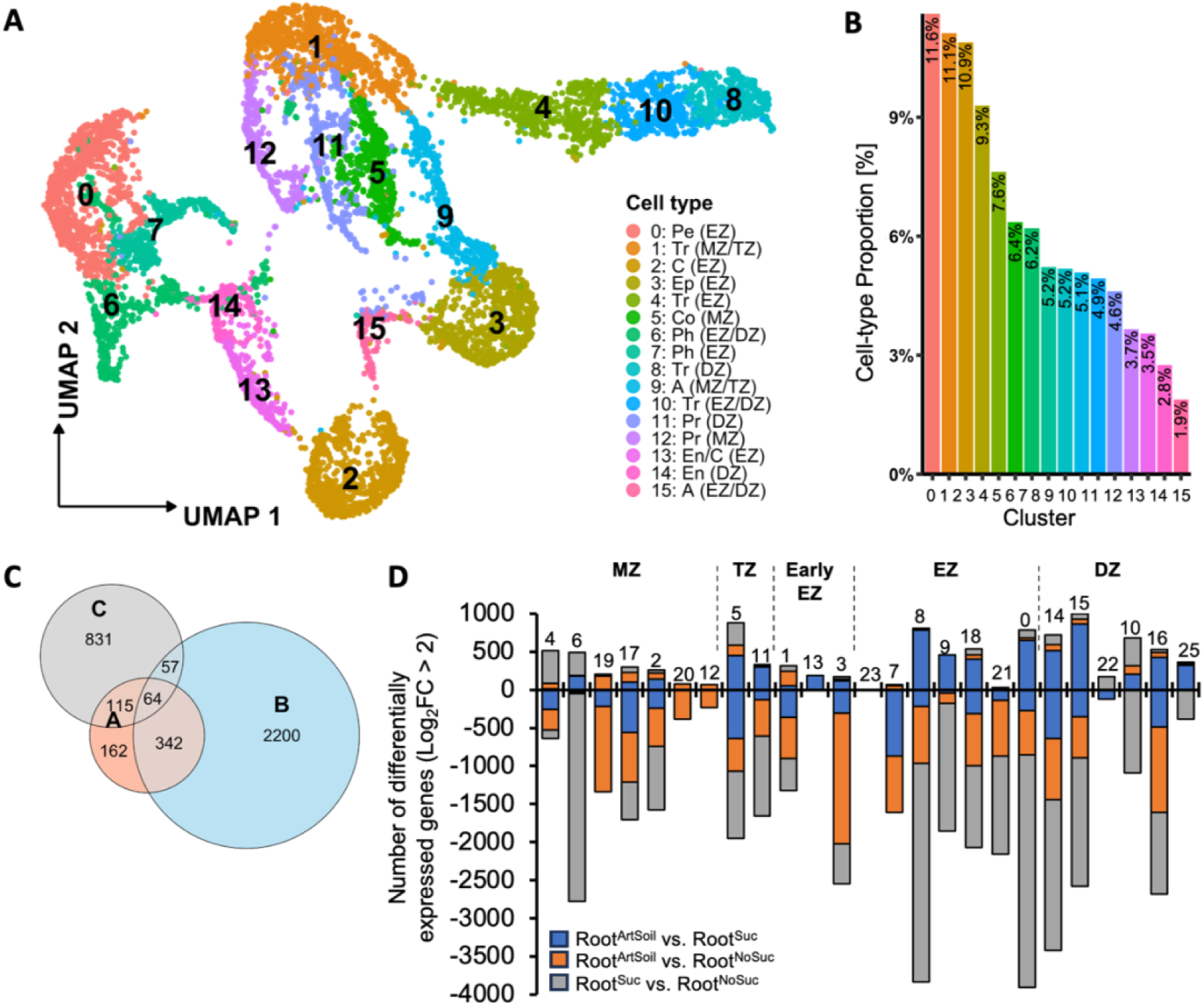
Single-cell transcriptomics of root from *A. thaliana* grown on ArtSoil. A) UMAP visualization of 8,000 single cells from 7-day-old *A. thaliana* roots grown on ArtSoil. Colors and numbers indicate clusters representing each cell type. Cell-types abbreviations: Pe- Pericycle, Tr- Trichoblast, C- Cortex, Ep- Epidermis, Co- Columella, Ph- Phloem, A- Atrichoblast, Pr- Procambium, En- Endodermis. Developmental zone abbreviations: MZ- Meristematic zone, EZ- Elongation zone, TZ- Transition zone, DZ- Differentiation zone. B) Relative abundance of annotated cell types among the 8,000 sequenced root cells. Bars represent the proportion of each cell type and developmental subtype identified within the dataset. Cluster number (X-axis) as indicated in (A). C) Venn Diagram depicting the number of differentially expressed genes (DEGs) with log_2_ fold-change >2 and adjusted p-value <0.05 from the comparison between the indicated single-cell transcriptomics datasets: A- ArtSoil vs ½ MS without sucrose supplementation, B- ArtSoil vs. ½ MS with sucrose supplementation, C- ½ MS with vs. without sucrose supplementation. D) Bar plot summarizing the number of DEGs up- and down-regulated by log_2_ fold-change >2 and adjusted p-value <0.05 in each indicated root cell type in ArtSoil. Labels above bars indicate root zones: MZ- meristematic zone, TZ- transition zone, EZ- elongation zone, DZ- differentiation zone. Numbers above each bar indicate cell type cluster (see Figure S2 for cluster annotation).

Single-cell transcriptomics of roots from ArtSoil (Root^ArtSoil^) was compared to published single-cell transcriptomics of roots of seedlings grown on sterile media supplemented with 1% sucrose (Root^Suc^)(Shahan et al., 2022) and roots of seedlings grown on sterile media without sucrose supplementation (Root^NoSuc^)(Wendrich et al., 2020). All three datasets were combined and clustered to identify the differentially expressed genes in each cluster (Figure S2A). A total of 6,204 genes were significantly upregulated in Root^ArtSoil^ compared to Root^Suc^ (adjusted p-value <0.05, log_2_FC >2), whereas 1,449 and 2,303 genes were significantly upregulated in Root^ArtSoil^ compared to Root^NoSuc^ and Root^Suc^ compared to Root^NoSuc^, respectively, indicating that sucrose inflated the number of genes induced in *A. thaliana* roots (Figure 5C). To inspect the extent to which the presence of microbes affects mRNA accumulation in a cell-type-specific manner, the number of up- and down-regulated genes was inspected for each cluster. The elongation and differentiation zones were the strongest affected by microbial presence on the root, as indicated by the largest variation in the number of DEGs (Root^ArtSoil^ vs. Root^Suc^/Root^NoSuc^) (Figure 5D). Along the root, the transcriptome of the meristematic zone (MZ) was least affected by the presence of microbes (Root^ArtSoil^ vs. Root^NoSuc^, number of upregulated genes: 600; Root^ArtSoil^ vs. Root^Suc^, number of upregulated genes: 447).

Studies using synthetic communities and analyses on bulk root tissues demonstrated that immunomodulation of plant defense activation by microbial community maintains plant growth homeostasis (Colaianni et al., 2021; Garrido-Oter et al., 2018; Ma et al., 2021; Teixeira et al., 2021). To inspect immunomodulation at the cellular level, defense signaling in Root^ArtSoil^ was inspected. Typical immunity pattern-recognition receptor genes were not highly induced in Root^ArtSoil^ (Figure 6A), in good accordance with microbial suppression of defense activation to maintain growth homeostasis. Root^ArtSoil^ retained a basal level of immune surveillance in the root (Zhou et al., 2020) as indicated by the co-expression of *PEPR1*, co-receptor *BAK1,* and ligands *PROPEPs* (Bartels et al., 2013; Krol et al., 2010; Yamaguchi et al., 2010, 2006) in the atrichoblast in the DZ (Figure 6A). *PROPEP6* was highly induced in the endodermis in the transition zone whereas *PROPEP4* was predominantly expressed in the MZ despite the lack of their corresponding receptors, suggesting a role beyond immune signaling, possible mobile peptide or yet to be discovered receptor. Notably, *PEPR1* was highly induced in almost all cell types in Root^Suc^, (Figure S2B) indicating sugar-induced defense activation, which may present artefacts in plant-microbe interaction studies using axenic media with sucrose supplementation.

**Figure 6:**
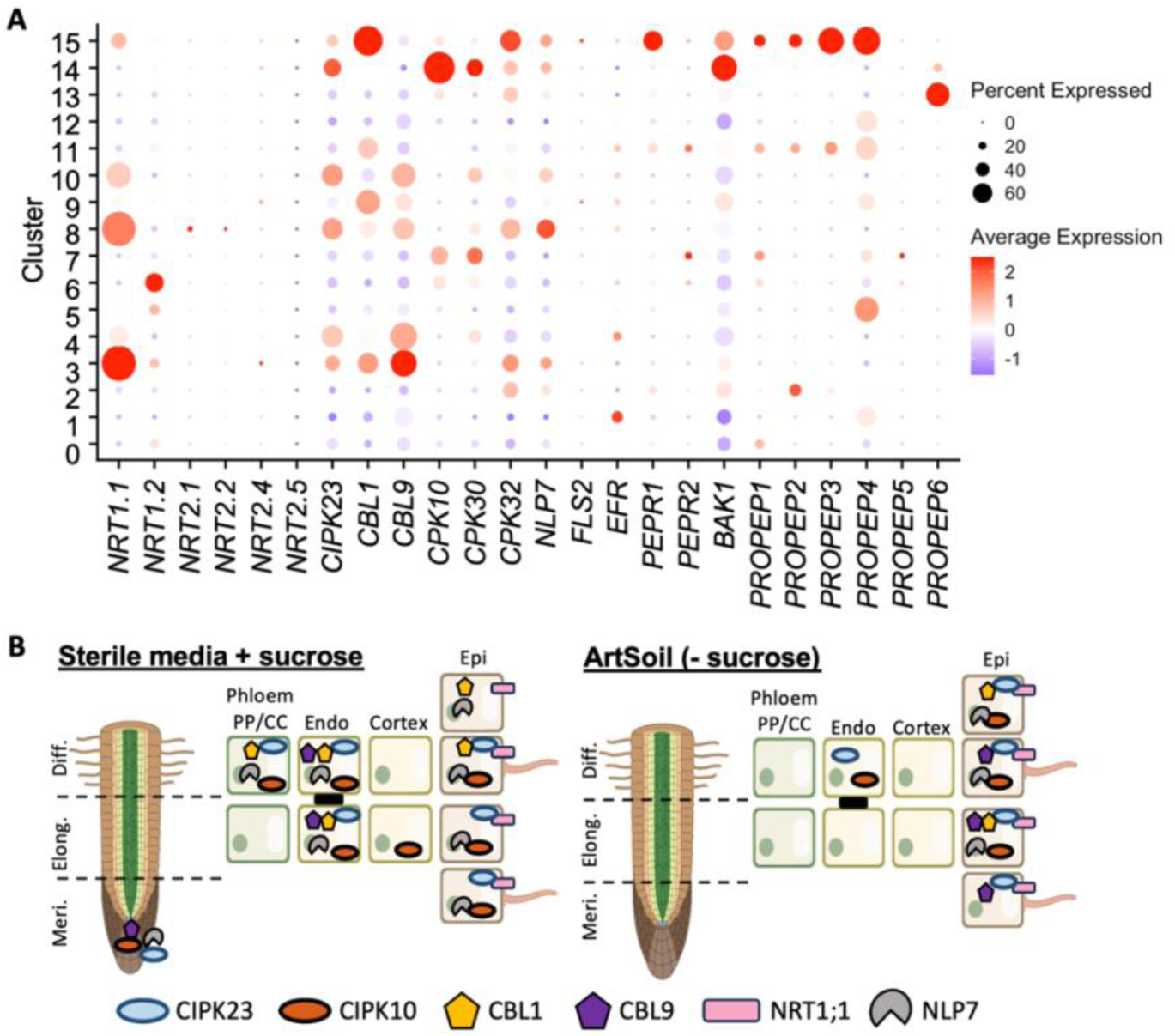
Microbiota-induced cell-type specificity in immune and nitrogen signaling. A) A dot plot describing the percentage of each cell type expressing the indicated genes in roots of plants grown in ArtSoil. Cell type represented by cluster numbers described in UMAP in Figure 5A. Sizes of dots indicate percentage of cells expressing the indicated genes, colors of dots indicate average expression of indicated genes. B) Cartoon depiction summarizing the presence of mRNA transcripts of five representative nitrogen uptake signaling components in roots grown on ArtSoil or media with sucrose supplementation (Shahan et al., 2022). Meri.- meristematic zone, Elong.- elongation zone, Diff.- differentiation zone, Endo- endodermis, Epi- epidermis.

The microbiota influences plant nutrient status and has been implicated to promote plant growth via nitrogen uptake (Calvo et al., 2019; Hernández-Reyes et al., 2022; Kechid et al., 2013; Mantelin, 2003). In nodulating plants, initial interaction between host and nitrogen-fixing bacteria occurs at the root hair epidermis (Oldroyd et al., 2011). To investigate whether microbiota-mediated nitrogen uptake could be cell-type specific in *A. thaliana*, the nitrogen signaling pathway was traced in Root^ArtSoil^. Among the nitrate transporters, only *NRT1.1* was upregulated in the root, specifically in the epidermal cells, i.e., the trichoblast and atrichoblast in the EZ and DZ. Following nitrate perception, a signaling cascade involving CIPK23 and its interacting partners CBL1 and CBL9 activates the master regulator NLP7 transcription factor (Huang et al., 1996; Liu et al., 2017; K.-H. Liu et al., 2022; Ródenas and Vert, 2021). *CIPK23* and *CBL1* were co-expressed with *NRT1.1* in the atrichoblast in the DZ, whereas *CIPK23* and *CBL9* were co-expressed in the trichoblast in the DZ (Figure 6A). Both *CBL1* and *CBL9* co-expressed with *CIPK23* and *NRT1.1* only in the atrichoblast in the EZ, indicating cell-type-specificity for *CBL1/9* interaction with *CIPK23* for the activation of *NRT1.1*. The master regulator *NLP7* was consistently co-expressed with *CIPK32,* but not with *CIPK10* and *CIPK30,* despite induction of the latter two genes in other cell types. Intriguingly, the co-expression of the signaling components in Root^ArtSoil^ indicates that the nitrogen signaling pathway is confined to the trichoblast in DZ and the atrichoblast in the EZ and DZ (Figure 6B). Tracing of the same nitrogen signaling module in Root^Suc^ showed non-cell-type-specific induction of *NRT1.1, CIPK23, CBL1, CBL9, NLP7*, and *CIPK32* (Figure 6B). Taken together, single-cell transcriptomics of roots from *A. thaliana* grown on ArtSoil revealed cell-type specificity in microbiota-induced immune and nitrogen signaling transcriptional reprogramming, and a clear distinction from Root^Suc^, possibly due to artefacts introduced by the presence of sucrose in the growth media.

### ArtSoil accommodates various aqueous soil extracts

To determine if ArtSoil is compatible with ASEs of other soils, four ASEs derived from native soils ArtSoil^CAS^, Reijerskamp (ArtSoil^REI^), Oranjewoud (ArtSoil^ORJ^) and Aijen (ArtSoil^AI^); and two commercial potting soils (i.e., soil enriched with macronutrients), Bedding Plant substrate (ArtSoil^BP^, https://klasmann-deilmann.com/de/anwendungsfelder/beet-und-balkonpflanzen) and Düsovit (ArtSoil^DUS^, https://www.duesovit.de/blumenerde.htm), were tested. Four-week-old plants grown on ArtSoil^CAS^, ArtSoil^BP^, and ArtSoil^DUS^ were significantly larger compared to plants grown on ½ MS without sucrose supplementation, whereas the shoot fresh mass of plants grown on ArtSoil^AI^, ArtSoil^ORJ^, and ArtSoil^REI^ was not significantly different from plants grown on ½ MS without sucrose supplementation (Figure 7A). To determine if soil nutrient content could contribute to possible phenotypic differences of plants grown on ArtSoil, the nutrient contents of the ASEs were determined via inductively coupled plasma mass spectrometry (ICP-MS, Figure 7B). Since the foundational media for ArtSoil (i.e., ½ MS) provides the essential nutrients for plant growth, we postulate that either the soil microbiota or non-essential nutrients carried over from the ASE may explain differences in plant growth (Anderson, 1956; Hewitt, 1951; Kumar et al., 2021). Of the eleven micronutrients carried over from the ASE, linear regression analysis indicated that strontium and platinum are significantly correlated with the shoot fresh mass of plants grown on ArtSoil (Figure S3). Ionomics on 1135 *A. thaliana* accessions revealed that strontium levels are positively correlated with calcium^68^, and early reports showed that strontium can be used as a substitute for the essential nutrient calcium to promote plant growth^69^. Although it cannot be excluded that strontium introduced from ASE could promote plant growth in ArtSoil^DUS^ and ArtSoil^BP^, its alternative, calcium supplied via ½ MS, is unlikely to be limiting in ArtSoil. All soil samples have traces of platinum, which may not explain significant differences in plant growth promotion by the different ASEs. Therefore, to disentangle the effects of nutrients from microbiota on plants grown on ArtSoil, two-week-old plants grown on sterile ArtSoil (i.e., without fresh soil solution, Figure 1) were evaluated. We posit that younger plants are more sensitive to nutrient availability due to higher metabolic demand for vegetative growth. In the absence of active soil microbiota, shoot fresh mass from plants grown on the various ASE tested was significantly larger than plants grown in ½ MS without sucrose supplementation (Figure 7C). Variation in shoot fresh mass between plants grown on the different ASEs was also reduced. Taken together, plant growth assay on sterile ArtSoil indicated that plant growth phenotype is largely attributed to the presence of soil microbial community, although traces of nutrients carried over from ASE contributed to plant growth, as evidenced by higher shoot fresh mass of plants grown on the sterile ArtSoils compared to ½ MS without sucrose supplementation. In sum, by investigating the various ASEs, ArtSoil was able to uncouple plant growth promotion conferred by microbiota from nutrients, providing possibilities for future mechanistic investigations.

**Figure 7:**
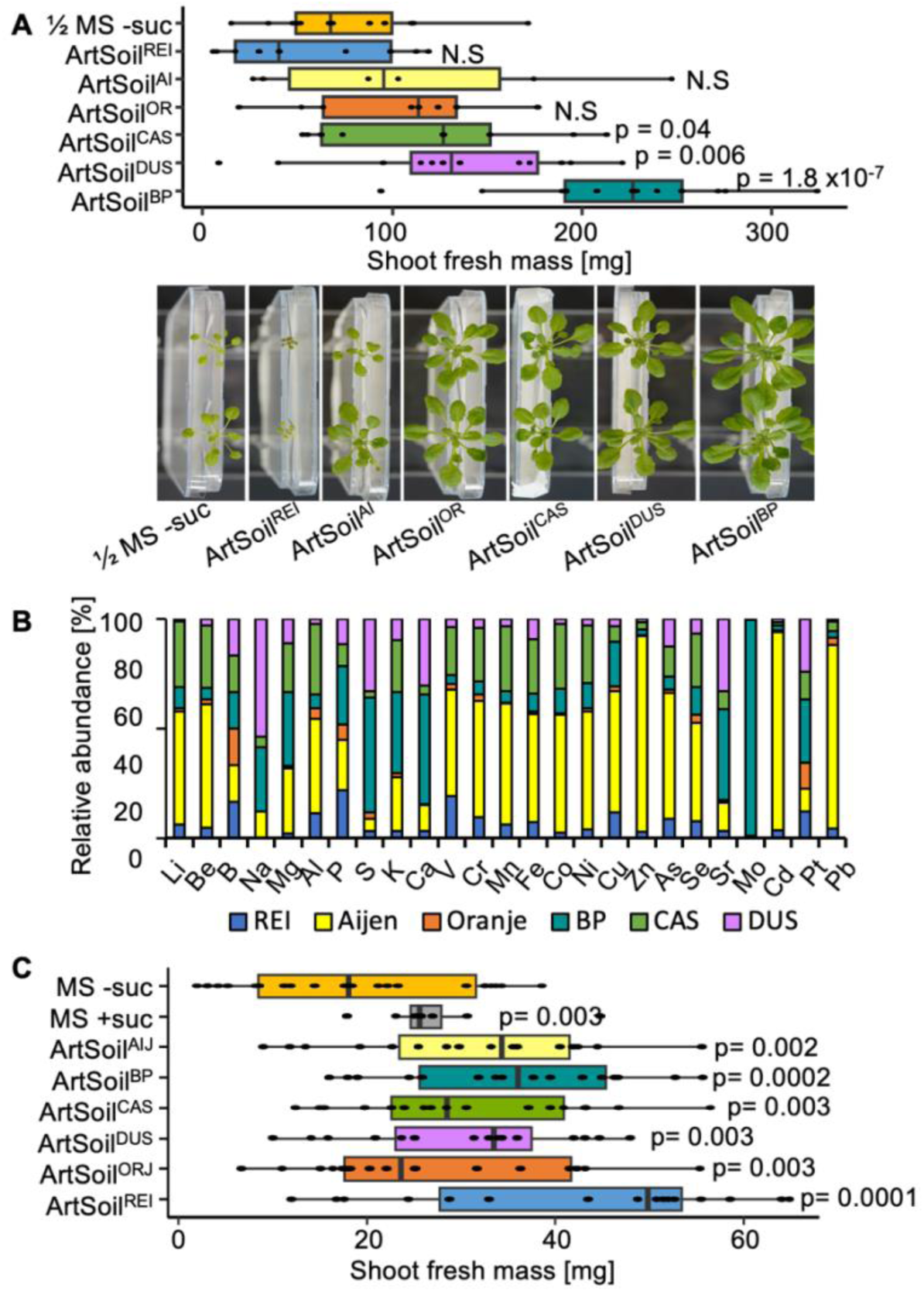
ASEs influence plant growth on ArtSoil. A) Top: Shoot fresh mass and phenotype and of 4-week-old *A. thaliana* grown on ArtSoil with indicated ASEs. Assay was repeated twice, N > 20. P value calculated via Student’s T-test against ½ MS without sucrose supplementation (½ MS -suc). N.S.- not significant. Bottom: phenotype of corresponding plants grown on indicated ASEs. B) Relative abundance of macro- and micronutrients in the respective soil used in ArtSoil as determined by inductively coupled plasma mass spectrometry (ICP-MS). Refer to Supplemental Table 1 for the absolute quantification of each nutrient. C) Shoot fresh mass of 2-week-old *A. thaliana* grown on sterile ArtSoil with indicated ASEs. Assay was repeated three, N > 20. P value calculated via Student’s T-test against ½ MS without sucrose supplementation (½ MS -suc). N.S.- not significant. The top and bottom boundaries of each boxplot indicate the 1^st^ and 3^rd^ quartiles, the center line indicates the median, and the whiskers represent 1.5× the interquartile range from the 1^st^ and 3^rd^ quartiles.

## Discussion

### Development of the ArtSoil

ArtSoil was developed to conserve soil microbial community and soil edaphic factors in the growth medium, thereby enabling plant cultivation under environments resembling the plant’s typical soil environment. ArtSoil was developed based on the criteria: i) readily available, ii) ease of use, and iii) customizable. ArtSoil requires materials and apparatus readily available in most laboratories and commonly used for plant growth, i.e., Petri dishes, growth media, and soil. ArtSoil does not require special tools or techniques for assembly and is similar to standard growth media preparation. Soil edaphic factors, i.e. pH, salinity, and nutrients are introduced into the growth medium via sterile ASE. The soil microbial population is subsequently introduced in controlled quantities using fresh soil. ArtSoil is compatible with most soils tested and is, therefore, customizable to the user’s choice of soil. Due to differences between soil, it is however important to either optimize the dilution of ASE and/or to test a set ASEs from different soils. Cultivation on ArtSoil does not differ from growing plants on other plate-based growth systems, e.g., in a growth chamber under controlled conditions. Since not all soils yielded optimal growth, users are encouraged to use a dilution series and test ASE from various soils by comparing phenotypes of plants grown on ArtSoil with the user’s soil of choice, ½ MS with and without sucrose, and/or ArtSoil with ASE from potting soil used in this study.

### ArtSoil for spatial characterization of the root

Though various plant growth systems have been developed for microbiota studies (Garrido-Oter et al., 2018; Korenblum et al., 2020; Kremer et al., 2021), a system that enables spatial profiling of the root microbiota has not been established prior to ArtSoil and CD-Rhizotron (Loo et al., 2024). Using ArtSoil and CD-Rhizotron, we previously profiled the spatial distribution of root microbiota and demonstrated that the spatial distribution of microbes along the longitudinal axis of *A. thaliana* root is not homogeneous (Loo et al., 2024). Technological advancements enable the investigation of plant responses at single-cell resolution (Denyer et al., 2019; Jean-Baptiste et al., 2019; Ryu et al., 2019). Single-cell sequencing technology has revealed heterogeneity in *A. thaliana* leaf transcription profiles, size and composition of local microbial populations in response to pathogen infection (Nobori et al., 2025; Saarenpää et al., 2023; Zhu et al., 2023). Similar to the leaf, *A. thaliana* root exhibits differential responses to microbial molecular patterns (Emonet et al., 2021; Zhou et al., 2020). Single-cell transcriptomics of *A. thaliana* roots have long been limited to sterile conditions due to the lack of suitable growth media, which hindered analyses in the presence of root microbiota, e.g., difficulties in harvesting young seedlings grown on soil without damaging roots, and in obtaining suitable protoplasts (free from soil particles) for subsequent cell sorting. ArtSoil enables single-cell resolution studies of microbiota-associated roots by facilitating large-scale cultivation of seedlings required to obtain sufficient protoplasts for single-cell sequencing. The cleaner roots derived plants grown on ArtSoil may also facilitate embedding and microsectioning for spatial transcriptomics, e.g., molecular cartography. Along with bacterial *in situ* hybridization, ArtSoil could be used for studying root responses to microbes at the single-cell level and examining transcriptomic differences between colonized and uncolonized cells. Notably, ArtSoil avoids the use of sucrose, and therefore will likely yield expression profiles that are more similar to those in native soil conditions, thereby enabling effective correlation of root gene expression profiles with microbiota colonization at single cell level.

An increasing number of studies have shown strong associations between root exudate composition and microbiota colonization (Koprivova and Kopriva, 2022; Korenblum et al., 2020; McLaughlin et al., 2023; Micallef et al., 2009). The state-of-the-art method for root exudate collection involves growing plants in nutrient media followed by transferring plants to distilled water or buffers. The nutrient media are sampled at specified time intervals and compared to a control (Döll et al., 2024; Oburger et al., 2013). Transferring plants from nutrient media to distilled water alters the exudation patterns due to increased analyte gradients, hypoosmotic shock, and possibly even plant cell lysis(Aulakh et al., 2001; Ayers and Thornton, 1968; Oburger et al., 2013). For instance, transferring maize seedlings from nutrient media to water alters the deposition rate of metabolites along the root zones and the rate of metabolite transfer across root growth zones (Walter et al., 2003). Root exudate collection using state-of-the-art methods thus masks metabolite heterogeneity along and across the root (Kranawetter et al., 2021; Loo et al., 2024; Moussaieff et al., 2013). ArtSoil combined with filter paper blotting on specific regions of the root (Neumann, Günter, n.d.) could allow spatiotemporal root exudate studies without the side effects associated with hydroponics systems and environmental shock during exudate sampling.

### Limitations of ArtSoil

Due to the strong preference of microbes for sugar (Estrela et al., 2021; Schäfer et al., 2023), ArtSoil is unsuitable for assays requiring media with sucrose supplementation, e.g., to rescue Arabidopsis mutant phenotypes. Sucrose availability in the media will result in microbial overgrowth/overcolonization of plants, potentially resulting in seedling death. ArtSoil may be unsuitable for research focused on nutrient depletion, unless the concentration of MS medium/nutrients is revised. Since ArtSoil encompasses nutrients from soil, ASE-derived nutrients in ArtSoil cannot be easily manipulated. An additional alternative could be to use soils with limited nutrient-of-interest and exogenously apply the nutrient-of-interest to the desired concentration, or use nutrient-depleting methods.

In conclusion, we introduce ArtSoil as an alternative growth medium that recapitulates the soil environment compared to the typically used conventional synthetic media. ArtSoil captures soil edaphic factors and microbiota that support plant growth without requiring sucrose supplementation. We show that plants grown on ArtSoil are influenced by three parameters, i.e., the soil used as ASE, the dilution of ASE, and whether the plants were grown in open or closed systems. We provide users with guidelines backed by empirical data to determine the most suitable ArtSoil parameters for the user’s experiment design. We demonstrated the application of ArtSoil in deciphering root-microbe transcriptome rewiring at the cellular level, and in spatial microbiota profiling (Loo et al., 2024). Finally, we recommend plant growth on ArtSoil essentially for all standard laboratory assays to maintain soil-like physiological conditions and to avoid artefacts/ side effects caused by the unphysiological external addition of sucrose. ArtSoil is not limited to growing *A. thaliana*. Rather, ArtSoil can be optimized for the growth of other plant species. In addition to characterizing root microbiota, ArtSoil can be used for a more comprehensive study of microbe-microbe-plant tripartite interactions where activities of pathogenic and/or beneficial microbes can be studied in the presence of the plant and its natural microbiota, in an ecologically-relevant growth environment. ArtSoil can also be used to pre-select the optimal synthetic community prior to use in other gnotobiotic systems.

## Materials and methods

### Plant growth conditions

Refer to supplementary Methods for detailed protocol of ArtSoil preparation. For all experiments, *Arabidopsis thaliana* wild-type Col-0 plants were used. Seeds were surface-sterilized in a 10% bleach solution supplemented with Triton X-100 for 10 minutes, followed by four washes with sterile water. Sterilized seeds were resuspended in sterile water and stratified in the dark at 4°C for at least three days prior to sowing. Plants were grown at 23°C with a 10-hour day/14-hour dark cycle and the light intensity of 25 or 120 μmol m^−2^ s^−1^, according to the experimental setup.

### Plant phenotyping

Total root length (TRL) was measured weekly for four weeks. At every measurement, images of the plates where seeds were sown were captured using a flatbed scanner (Epson Perfection V850 Pro). TRL was measured using the FIJI software, using the “freehand draw” tool. Roots were traced, and the length was measured using the “measure” tool. The scale was calibrated to centimeters. The TRL is the sum of the length of the main and lateral roots developed by a single plant. For shoot fresh mass, the shoots were cut at the hypocotyl using tissue scissors. Shoots were tapped dry before weighing using an analytical scale.

### Root protoplast isolation

Root protoplasts were isolated following a protocol adapted from (Birnbaum et al., 2003), optimized for single-cell RNA sequencing. Seven-day-old *A. thaliana* Col-0 seedlings grown on ArtSoil were harvested by cutting approximately 1.5 cm from the root tip with a razor blade. Roots were rinsed in protoplast buffer (0.6 M mannitol, 0.1 M KCl, 0.02 M MgCl₂, 0.02M CaCl₂ 0.08 M MES, and 0.1% BSA (Sigma-Aldrich), adjusted to pH 5.5 with 0.1 M Tris-HCl) for approximately five minutes to remove debris and surface microbes. The roots were transferred into 5 mL of freshly prepared enzyme solution containing 1.5% Cellulase R-10 and 0.1% Pectolyase (Duchefa Biochemie) dissolved in root protoplast buffer composed of 0.6 M mannitol, 0.1 M KCl, 0.02 M MgCl₂, 0.02 M CaCl₂, 0.08 M MES, and 0.1% BSA (Sigma-Aldrich), adjusted to pH 5.5 with 0.1 M Tris-HCl. Roots were digested at 26°C for 2 hours with gentle shaking (200 rpm) on an orbital shaker. After digestion, the protoplast suspension was passed through a 100 µm nylon mesh filter and rinsed with 1 - 5 mL of wash buffer. Cells were centrifuged for 10 minutes at 500 × g at 4°C. The supernatant was carefully removed, and the pellet was resuspended in 10 mL of wash buffer. This washing step was repeated once, and the final pellet was resuspended in ≤ 500 µL of the same buffer. Protoplasts were visually inspected under a light microscope, and, if necessary, residual debris was removed by an additional washing step. The cell suspension was passed through a 40 µm cell strainer (Corning). Protoplasts were stained with 4% Trypan blue and viable cells were quantified using a hemocytometer. The final concentration was adjusted to 500 - 1000 cells/µL.

### Single Cell Library Generation

A total of ∼8,000 single cells were generated on the 10X Chromium Controller system for single-cell droplet library generation utilizing the Chromium Single Cell 3’ GEM-X Reagent Kit v4 (10X Genomics, Pleasanton, CA, USA) according to manufacturer’s instructions. Sequencing was carried out on a NextSeq2000 system (Illumina Inc. San Diego, USA) with a mean sequencing depth of ∼25,000 reads/cell.

### Processing of 10X Genomics single cell data

Raw sequencing data was processed using the 10X Genomics CellRanger software (v9.0.1). Raw BCL-files were demultipexed and processed to Fastq-files using the CellRanger *mkfastq* pipeline. Alignment of reads to the *A. thaliana* TAIR10_59 genome build and UMI counting was performed via the CellRanger *count* pipieline to generate a gene-barcode matrix. Further analyses were carried out with the Seurat v5.1.0 R package (Butler et al., 2018; Hao et al., 2021; Stuart et al., 2019). Initial quality control consisted of removal of cells with fewer than 200 detected genes as well as removal of genes expressed in less than 3 cells. Cell doublets have been removed from the dataset using DoubletFinder v2.0 (McGinnis et al., 2019). Normalization has been carried out utilizing SCTransform. Dimensional reduction of the data set was achieved by Principal Component analysis (PCA) based on identified variable genes and subsequent UMAP embedding. The number of meaningful Principal Components (PC) was selected by ranking them according to the percentage of variance explained by each PC, plotting them in an “Elbow Plot” and manually determining the number of PCs that represent the majority of variance in the data set. Cells were clustered using the graph-based clustering approach implemented in Seurat. Markers defining each cluster as well as differential gene expression between different clusters were calculated using a Wilcoxon Rank Sum test which is implemented in Seurat.

### Soil ICP-MS analysis

Fresh soil samples were dried in a 60°C oven for 48 hours. The total mineral content was determined by inductively coupled plasma mass spectrometry (ICP-MS) with a 7700 system from Agilent (Waldbronn, Germany). To 100 mg homogenized soil, 500 µL of 30% HNO_3_ were added and let soak for 2-3 hours. Another 500 µL of 30% HNO_3_ were added and the samples were placed overnight in water bath at 65°C. Afterwards the samples were boiled at 95°C for 30 min. After cooling down the sample to room temperature 200 µL of 30% H_2_O_2_ were added and boiled again for 30 min at 95°C. To reach a final volume of about 10 mL 8.8 ml of water was. The samples were kept overnight by 4°C, afterwards particles were spun down at 7000 g, 4°C for 1 hour or at 3500 G for 3 hours. 0.8 mL of supernatant was mixed with 3.2 mL of 2% HNO_3._ The elemental concentration was determined using the Agilent 7700 ICP-MS following the manufacturer’s instructions.

### Root microbiota profiling

Total DNA was extracted using NuceloSpin Soil Mini kit for DNA from soil (Macherey-Nagel GmbH & Co. KG, Düren, Germany) following instructions from the manufacturers. DNA samples were eluted in 50 μL nuclease-free water and used for microbial community profiling. DNA samples were used in a two-step PCR amplification protocol using primers targeting the V4-V7 16S rRNA region (799F: AACMGGATTAGATACCCKG; 1192R: ACGTCATCCCCACCTTCC) using 2 U DFS-Taq DNA polymerase, 1× incomplete buffer (Bioron GmbH, Ludwigshafen, Germany), 2 mM MgCl2, 0.3% BSA, 0.2 mM dNTPs (Life technologies GmbH, Darmstadt, Germany) and 0.3 μM forward and reverse primers. PCR was performed using the following parameters: 94 °C/2 min, 94 °C/30 s, 55 °C/30 s, 72 °C/30 s, 72 °C/10 min for 25 cycles. PCR products were pooled, end-repaired, A-tailed, and ligated with Illumina adapters. Amplicon concentration was determined fluorescently (Quant-iT™ PicoGreen™, Invitrogen), and equivalent DNA amounts of each of the barcoded amplicons were pooled in one library. The final library concentration was estimated fluorescently (Quantus™ Fluorometer, Promega). Paired-end Illumina sequencing was performed in-house using the MiSeq sequencer and custom sequencing primers at the Max Planck Institute for Plant Breeding Research.

## Acknowledgement

This research has been supported by the Deutsche Forschungsgemeinschaft (DFG, German Research Foundation) under Germanýs Excellence Strategy – EXC-2048/1 – project ID 390686111, DFG project ID 391465903/GRK 2466 and the Alexander von Humboldt Foundation (AvH professorship to Wolf Frommer). Computational infrastructure and support were provided by the Centre for Information and Media Technology at Heinrich Heine University Düsseldorf. We thank the lab of Bart Thomma for contributing soil samples and Wolf Frommer for scientific discussions.

## Author contributions

Conception: EL; Writing: EL, VK, HA; Plant phenotyping: HA, TG, EL; Microbiota: PD, EL; Single-cell transcriptomics: VK, EL, TL; Soil analysis: SM, EL.

## Declaration of interests

The authors declare no competing interests.

## Legends for supporting information

Figure S1: Time-series quantification of total root length of *A. thaliana* grown under optimal or sub-optimal light conditions on ArtSoil, ½ MS with or without sucrose supplementation.

Figure S2: Transcriptomics analyses for Root^ArtSoil^, Root^Suc^, and Root^NoSuc^.

Figure S3: Linear regression analysis on the influence of individual soil nutrients on *A. thaliana* shoot fresh mass.

Table S1: Soil elemental analysis for the amount (mg/kg) of each nutrient found in respective soil sample.

Methods S1: Detailed protocol for the preparation of ArtSoil.

## Notes

### Competing Interest Statement

The authors have declared no competing interest.

