## Supplemental materials for "Artificial Soil (ArtSoil): recreating soil conditions in synthetic plant growth media"

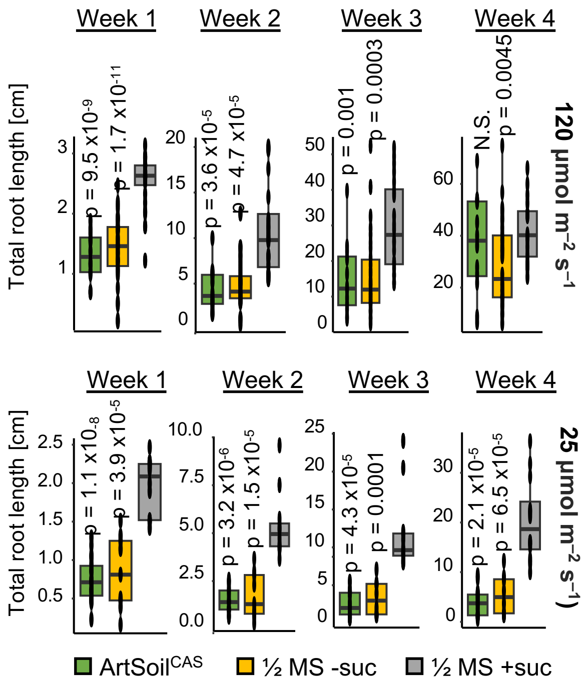


**Supplementary Figure 1: Time-series quantification of total root length of *A. thaliana* grown under optimal or sub-optimal light conditions on ArtSoil, ½ MS with or without sucrose supplementation.** Total root length of 4-week of *A. thaliana* grown on ½ MS with or without sucrose, and ArtSoil in optimal and sub-optimal light conditions. All assays were repeated three times, N > 15. P values were calculated via Student's T-test against ½ MS with sucrose supplementation (½ MS +suc). The top and bottom boundaries of each boxplot indicate the 1st and 3rd quartiles, the center line indicates the median, and the whiskers represent 1.5× the interquartile range from the 1st and 3rd quartiles.

A

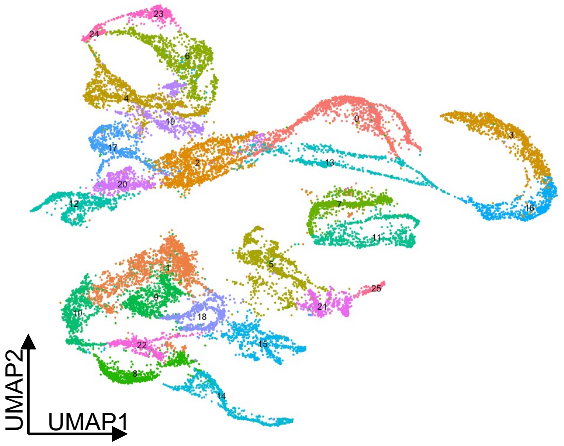


| Cluster | Cell type | Zone |
|---|---|---|
| 0 | Atrichoblast | Late EZ |
| 1 | Xylem, phloem pericycle | Early EZ |
| 2 | Epidermis | MZ |
| 3 | Trichoblast | Early EZ |
| 4 | Detaches cell | MZ |
| 5 | Endodermis | TZ |
| 6 | LRC | MZ |
| 7 | Cortex | EZ |
| 8 | Procambium | Late EZ |
| 9 | Xylem pole pericycle | Late EZ |
| 10 | Procambium | DZ |
| 11 | Cortex | TZ |
| 12 | Endodermis | Late MZ |
| 13 | Trichoblast | Early EZ |
| 14 | Protoxylem | DZ |
| 15 | Phloem companion cell | DZ |
| 16 | Trichoblast | DZ |
| 17 | Initial cells | MZ |
| 18 | Xylem pole pericycle | Late EZ |
| 19 | Columella | MZ |
| 20 | Atrichoblast | MS |
| 21 | Endodermis | DZ |
| 22 | Phloem companion cell, procambium | DZ |
| 23 | Phloem sieve element | EZ |
| 24 | LRC detached cells | MZ |
| 25 | Endodermis | DZ |

B

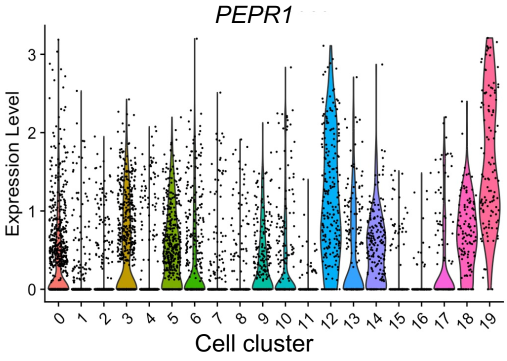


**Fig S2: Transcriptomics analyses for Root$^{ArtSoil}$, Root$^{Suc}$, and Root$^{NoSuc}$**

A) Merged and reclustering of single-cell transcriptomics of *A. thaliana* seedlings grown on media supplemented with sucrose (Shahan et al., 2022), without sucrose (Wendrich et al., 2020), and ArtSoil. Cell identity and root zones of each cluster listed in table on the right. MZ- meristematic zone, EZ- elongation zone, TZ- transition zone, DZ- differentiation zone, LRC- lateral root cap.

B) mRNA transcript level of *PEPR1* each cell type in Root$^{Suc}$ (Shahan et al., 2022)

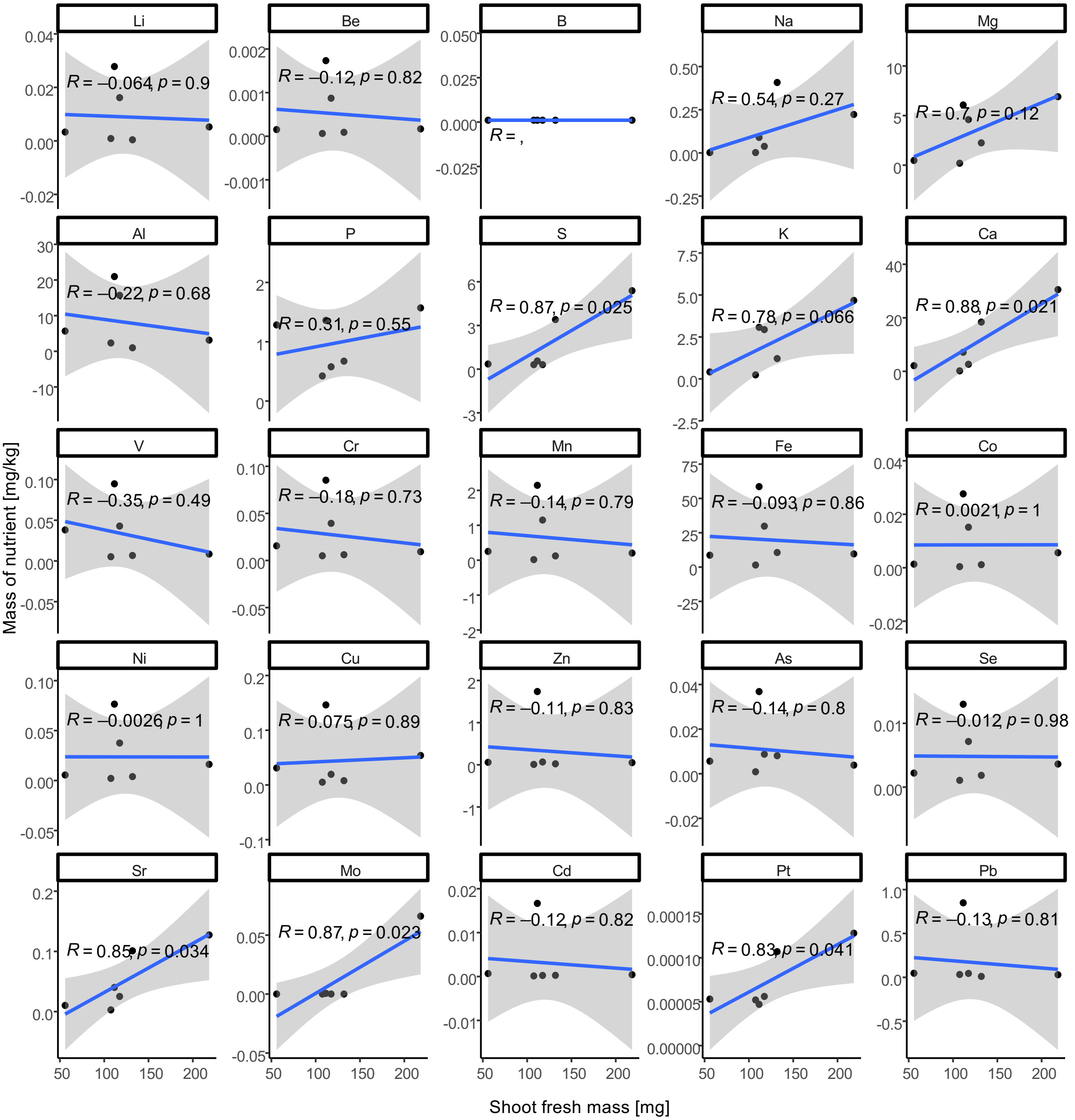


**Figure S3: Linear regression analysis on the influence of individual soil nutrients on *A. thaliana* shoot fresh mass.**

Supplemental Table 1: Soil elemental analysis for the amount (mg/kg) of each nutrient found in respective soil sample.

| | Li | Be | B | Na | Mg | Al | P | S | K | Ca | V | Cr | Mn | Fe | Co | Ni | Cu | Zn | As | Se | Sr | Mo | Cd | Pt | Pb |
|---|---|---|---|---|---|---|---|---|---|---|---|---|---|---|---|---|---|---|---|---|---|---|---|---|---|
| Reijeskamp | 0.0033 | 0.0002 | 0.0010 | 0.0022 | 0.4758 | 5.6727 | 1.2841 | 0.3526 | 0.4149 | 2.1112 | 0.0381 | 0.0155 | 0.2516 | 8.6555 | 0.0013 | 0.0056 | 0.0310 | 0.0588 | 0.0056 | 0.0022 | 0.0102 | 0.0000 | 0.0007 | 0.0001 | 0.0466 |
| Aijen | 0.0278 | 0.0017 | 0.0010 | 0.0898 | 6.0716 | 20.9599 | 1.3611 | 0.5609 | 3.0720 | 7.1224 | 0.0947 | 0.0852 | 2.1456 | 58.6169 | 0.0276 | 0.0765 | 0.1463 | 1.7379 | 0.0368 | 0.0130 | 0.0402 | 0.0006 | 0.0167 | 0.0000 | 0.8494 |
| Oranjewound | 0.0009 | 0.0001 | 0.0010 | 0.0022 | 0.1958 | 2.3423 | 0.4229 | 0.3091 | 0.2382 | 0.1725 | 0.0052 | 0.0050 | 0.0170 | 1.4950 | 0.0004 | 0.0021 | 0.0051 | 0.0087 | 0.0009 | 0.0011 | 0.0028 | 0.0000 | 0.0001 | 0.0001 | 0.0325 |
| BP-substrate | 0.0052 | 0.0002 | 0.0010 | 0.2226 | 6.9043 | 3.1354 | 1.5709 | 5.3862 | 4.6766 | 30.5240 | 0.0085 | 0.0093 | 0.2036 | 9.5876 | 0.0056 | 0.0162 | 0.0539 | 0.0507 | 0.0038 | 0.0036 | 0.1274 | 0.0661 | 0.0004 | 0.0001 | 0.0294 |
| Cologne Agricultural Soil | 0.0162 | 0.0009 | 0.0010 | 0.0381 | 4.5933 | 15.6914 | 0.5774 | 0.3091 | 2.9371 | 2.6382 | 0.0429 | 0.0394 | 1.1498 | 29.7945 | 0.0152 | 0.0376 | 0.0195 | 0.0666 | 0.0087 | 0.0072 | 0.0251 | 0.0000 | 0.0002 | 0.0001 | 0.0447 |
| Düsovit | 0.0004 | 0.0001 | 0.0010 | 0.4079 | 2.2487 | 0.9856 | 0.6730 | 3.4047 | 1.2109 | 18.3928 | 0.0067 | 0.0061 | 0.1228 | 10.6140 | 0.0011 | 0.0040 | 0.0079 | 0.0232 | 0.0081 | 0.0018 | 0.1010 | 0.0000 | 0.0003 | 0.0001 | 0.0106 |

**Method S1: Detailed protocol**

***Box 1: A.thaliana seed preparation (Figure 1)***

Timing: 3-5 days

Reagents and materials

- *A. thaliana* seeds
- Commercial bleach (DanKlorix)
  Caution: Bleach is corrosive. Handle with appropriate safety measurements.
- Triton X-100 (Sigma-Aldrich, CAS No.: 9002-93-1)
  Caution: Triton X-100 is hazardous and harmful if swallowed and can cause eye damage/irritation. Handle with using appropriate safety measurements.
- Autoclaved deionized distilled water ($ddH_2O$)
- 80% (volume/volume) ethanol (Sigma-Aldrich, CAS No.: 30899-19-5)
  Caution: Ethanol is flammable. Handle with appropriate safety measurements.

Equipment

- 1.5 mL microcentrifuge tube (Sarstedt, Cat. No. 72.690.001)
- 50 mL centrifuge tube (Sarstedt, Cat. No. 62.547.254)
- Micropipette and tips
- VorteX-Genie 2 (Scientific Industries Inc., SKU: SI-0256)
- Vortex adapter- Microtube foam insert (Scientific Industries Inc., SKU: 504-0234-00)
- Benchtop centrifuge
- Tea sieve

Post-harvest seed cleaning

Timing: 30 min

1. Place mature and dried *A. thaliana* siliques into a clean tea sieve. Using a clean finger, gently rub the siliques against the sieve to release the seeds.
2. Shake by tapping the side of the sieve to filter the seeds onto a piece of clean paper. Since the diameter of seeds is smaller than the pore size of the sieve, the seeds will fall through the sieve onto the paper, whereas empty siliques and other debris will remain in the sieve. Discard the debris.
3. Place the fall-through seeds back into the sieve and repeat step 2 until seeds are free of debris.

Preparation of seed sterilization solution

Timing: 5 min

4. Add 15 mL of commercial bleach and 10 µL of Triton X-100 into a clean 50 mL tube. Top up to a final volume of 50 mL with $ddH_2O$. Shake well. Seed sterilization solution can be stored at room temperature for several months.

Seed sterilization

Timing: 30 min

5. Fill a clean 1.5 mL microcentrifuge tube with an amount of *A. thaliana* seeds according to your intended application.

6. Add 500 µL of seed sterilization solution to the seeds. Incubate with shaking for 15-20 minutes. We use a vortex with a microcentrifuge tube adaptor. However, any device that could agitate the seeds would work, e.g. a rotary shaker.
   Caution: Do not incubate seeds in seed sterilization solution over 1 hour. Keep an eye on the colour of the seeds. Do not use the seeds if the colour is bleached.
7. Spin down briefly (3-5 seconds) using a benchtop centrifuge to sediment the seeds.
8. Carefully remove the seed sterilization solution using a micropipette.
   Critical: The seeds may clump when treated with seed sterilization solution. If seeds clump, vortex until the clump is dissolved.
9. Add 500 µL of 80% ethanol. Vortex for 1-3 seconds, followed by a brief spin-down to pellet the seeds. Carefully remove the ethanol using a micropipette.
   Critical step: The seeds may clump when treated with ethanol. If seeds clump, vortex until the clump is dissolved.
10. Add 750 µL of sterile dd$H_2O$ to rinse the seeds. Vortex for 1-3 seconds. Pellet the seeds via a brief spin-down. Carefully remove the dd$H_2O$ using a micropipette.
11. Repeat the rinsing step (Box 1 step 10) three times, each time with fresh sterile dd$H_2O$.
12. After the final rinse, discard spent dd$H_2O$ and add 1 mL of fresh sterile dd$H_2O$.
13. Wrap the tube in aluminum foil and keep in 4°C for 3-5 days.

*Troubleshooting*

***Box 2: Preparation of half-strength Murashige-Skoog media (½ MS)***

Timing: 1.5 hrs (including autoclave)

Reagents

- Murashige & Skoog Medium with basal salt including MES buffer (MS medium) (Duchefa Biochemie, Cat. No.: M0254.0050)
  Critical step: Make sure to use MS medium without added sugar
- Agar-agar for plant (Roth, CAS No.: 9002-18-0)
- dd$H_2O$
- 1 M Potassium hydroxide (KOH)

Equipment

- 1 L media bottle with cap
- Weighing scale
- pH meter

Preparation

1. Add 2.4 g of MS medium and 10 g of agar into a media bottle. Add 800 mL dd$H_2O$.
   Critical: 1L of MS media produces 20 plates (12 cm x 12 cm Petri dishes of ArtSoil)
2. Adjust the pH of MS media to 5.7 with 1M KOH.
   Critical step: Add KOH drop-by-drop. Typically, only several drops of 1M KOH are required to achieve a pH of 5.7.
3. Top up the media to 1 L with dd$H_2O$.
4. Autoclave at 121°C for 15 mins.
   Critical step: Make sure to autoclave the medium shortly after its preparation to avoid contamination. Keep media at 4°C for longer storage before autoclaving.

Pause Point: Autoclaved ½ MS can be stored at room temperature for several weeks.

***Part 1: Preparation of 10% autoclaved soil solution (10% AS)***
Timing: 3 days
Biological materials
*A.thaliana* seeds.
Fresh soil

Critical step: Keep soil at 4°C in a sealed bag or container. We recommend airing the soil at RT once a month to prevent buildup of excessive moisture that may lead to algae/fungal bloom.

Equipment

- 50 mL centrifuge tubes (Sarstedt, Cat. No. 62.547.254)
- 100 mL media bottle with cap
- Infrared thermometer
- Micropipette and tips
- Weighing scale
- Sieve
- VorteX-Genie 2 (Scientific Industries Inc., SKU: SI-0256)
- 12 cm x 12 cm square Petri dishes (Greiner, Cat. No. 688161)
- Micropore tape (3M, Cat. No. 1530-0)
- Biosafety cabinet (Heraeus LaminAir HB 2448)
- Plant growth chamber

Procedure

1. For 50 mL of 10% AS, add 5 g of fresh soil into a clean 100 mL media bottle. Add 50 mL of $ddH_2O$ into the media bottle. Shake the bottle thoroughly to ensure soil clumps, if present, are dissolved. 50 mL of 10% AS can be used to produce up to 4 L of ArtSoil media ($10^{-4}$ dilution) or 80 Petri dishes (12 cm x 12 cm square Petri dishes).
   Critical step: Stir to homogenize the soil before use. If the soil has a high pebble content, we recommend sieving the soil. Scoop the soil with a clean spatula.
   Critical step: We do not recommend preparing lower than 10 mL 10% AS (i.e. 1 g of soil) since using a low amount of soil may bias towards only high-abundance microbes.
2. Autoclave at 121°C for 15 mins for three consecutive times. Allow the 10% AS to sit overnight (>16 hours) in between each autoclave session. After each autoclave session, shake the bottle to fully resuspend the soil.
   Critical step: Make sure the 10% AS rests overnight before the next autoclave step.
3. After the final autoclave, allow 10% AS to cool to the touch before use.

Pause Point: 10% AS can be prepared in bulk and stored at room temperature for several weeks.

***Part 2: Preparation of Aqueous Soil Extract (ASE)***
Timing: 15 mins.

Critical: ASE and ArtSoil need to be prepared freshly before every use.

Preparation of 50% fresh soil solution (50% FS)

4. Prepare the 50% (weight/volume) FS by mixing 3 g fresh soil and 6 mL 10% AS in a clean 50 mL centrifuge tube

   Critical step: Thoroughly shake 10% AS before use to ensure homogenous distribution.

   Critical step: Prepare 50% soil suspension in a biosafety cabinet under sterile conditions.

5. Vortex the 50% FS vigorously and let stand at room temperature for 10-15 minutes.

Preparation of ASE dilution ($10^{-2}$ dilution)

6. In a clean 50 mL centrifuge tube, dilute 50% FS using 10% AS to achieve 100x dilution. For example, to prepare 30 mL of ASE $10^{-2}$ dilution, use 300 µL of 50% FS and top up to 30 mL with 10% AS.

   Critical step: Thoroughly shake the 10% AS to ensure sediments are resuspended before use.

   Critical step: Prepare Dilution 1 in a biosafety cabinet under sterile conditions.

***Part 3: Preparation of ArtSoil (with $10^{-4}$ dilution)***

Timing: 15 mins.

See Box 3 for the preparation of open system ArtSoil.

7. Cool ½ MS to about 40°C. Use an infrared thermometer for accurate temperature measurement.

   Critical step: If ½ MS has solidified, melt the media by microwaving and incubate in 40°C water bath until use.

8. Use a 1:100 ratio of ASE $10^{-2}$ dilution: ½ MS media, i.e., use 1 mL of ASE $10^{-2}$ dilution for every 100 mL of ½ MS media. Gently swirl the media bottle to ensure thorough mixing whilst avoiding formation of bubbles.

   Critical step: Thoroughly shake ASE $10^{-2}$ Dilution to ensure sediments are resuspended before use.

   Critical step: Agar solidifies at 36°C ± 2°C. Work quickly but carefully to prepare the plates before the agar sets. If longer preparation time is required, it is possible to reduce the agar to 0.8% (weight/volume).

   Critical step: Prepare ArtSoil in a biosafety cabinet under sterile conditions.

   Critical step: Steps 6-8 describe the protocol to achieve $10^{-4}$ dilution of ArtSoil. Users should test a series of dilutions (Figure 2) to determine the most suitable dilution for the user's soil of choice.

9. Measure 50 mL of ArtSoil media using a sterile 50 mL centrifuge tube. Gently pour the media into a 12 cm x 12 cm square Petri dish.

   Critical step: 50 mL media per 12 cm x 12 cm square petri dish is suitable for assays that last 4-5 weeks. Assays that require longer plant growth should adjust the volume of media accordingly and/or use a different growth vessel.

   Critical step: Pour media gently to avoid air bubbles. If bubbles form, pop them using a sterile pipette tip or push them to the edges of the Petri dish.

10. Allow the media to set in a clean bench, typically 15-20 mins.

    Critical step: Keep the lid of the plate slightly ajar to prevent condensation on the lid.

*Troubleshooting*

***Part 4: Seed sowing***
Timing: 30 mins.

11. Aspirate the sterilized seeds (Box 1) using a sterile 1000 μL micropipette.
12. Carefully remove the pipette tip from the micropipette without dispensing the seeds. Hold the pipette tip vertically to allow the seeds to settle at the tip of the pipette tip. If seeds do not sediment at the tip, gently tap the tip to facilitate seed descent.
13. Gently tap the pipette tip on the surface of the ArtSoil media to dispense the seeds. Make sure to dispense only one seed per tap.
    Critical step: The number of seeds sown per tap can be adjusted based on the amount of liquid the seeds are resuspended in, the angle of the pipette tip against the media, and how quick each tap is.
14. Seal the plates using micropore tape.
    Critical step: Make sure the petri dish is sealed properly, especially at the corners of the Petri dish.
15. Incubate the plates vertically in a growth chamber for up to 5 weeks.
    Critical step: Adjust growth condition according to your experiment requirements.
    Critical: Colony formation on ArtSoil is normal and is not indicative of contamination (Figure 6A). The size and colour of the colonies differ based on the soil used. A telltale of contamination is typically black fungi or retarded plant growth when plants come in contact with the fungi.

*Troubleshooting*

***Box 3: Preparation of open system ArtSoil***
Timing: 1 hr.
The open system has two outlets on one side (i.e., the top) of the plate to allow the plant shoots to emerge (Figure 2, Figure 4). Users are recommended to refer to Nagel *et al.,* 2020 for more information on the construction of an open system.

Reagents

- ArtSoil media: preparation as previously described in Parts 1-3.

Equipment

- 12 cm x 12 cm square Petri dishes (Greiner, Cat. No. 688161)
- Pipette and 1 mL tips
- Tape
- Aluminum foil or dark paper
- Biosafety cabinet (Heraeus LaminAir HB 2448)
- Handheld borer with 3 mm diameter head (Dremel 8200).
- Growth chamber

Procedure

1. Measure 2 cm from each corner of one side of a 12 cm x 12 cm square Petri dish and mark using a permanent marker (Figure M1C).
2. Tape the lip to the body to hold the Petri dish close.
3. Using a handheld borer (3 mm diameter, Figure M1A), drill through the body of the petri dish to make two outlets for plant shoots.

4. Critical step: Make sure you drill only through the body of the Petri dish, not the lid (Figure M1B).
5. Temporarily seal the holes with tape to prevent media leakage.
6. Remove the tape that held the lid to the body of the Petri dish. Measure 100 mL of ArtSoil media and gently pour the media into the Petri dish.
   Critical step: Make sure the holes are filled completely with ASE. Partially filled holes may lead to seeds falling off the agar.
7. Once the media solidified, remove the tapes that were used to cover the holes.
8. Sow one seed directly on each drilled opening of the plate using a pipette as described in Part 4.
   Critical step: It is not recommended to expose the Petri dish unnecessarily to the surrounding atmosphere. If possible, conduct all steps in a biosafety chamber.
9. Wrap the Petri dish with aluminum foil or a black piece of paper prior to incubation in the plant growth chamber.

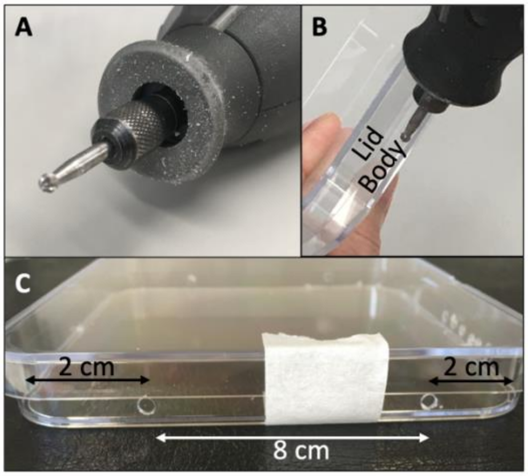


**Figure M1: Construction of open-system ArtSoil.**

A) Handheld borer with 3 mm diameter head. The model used in this work was Dremel 8200 but any device that serves the same purpose would work.
B) Drill into the body of a square petri dish.
C) Measurements and layout of holes on a square petri dish. Notice the tape used to hold the lid and body together while drilling.

Troubleshooting

| Step | Problem | Possible reason | Solution |
|---|---|---|---|
| Box 1 Step 12 | Seeds clump together | Seed sterilization solution or ethanol was not rinsed of properly. | Repeat the rinsing step (Box 1, step 10) until seeds do not clump |
| | | High seed density in the microcentrifuge tube. | Reduce the amount of seed per microcentrifuge tube. |
| Part 3 Step 10 | ArtSoil components are not homogeneously distributed | ArtSoil media solidified before adding ASE or pouring. | Gently shake the ArtSoil media after adding soil inoculum. |
| | | | Prevent the medium from cooling below 40°C. Work quickly. |
| | | | Reduce the agar to 0.8% (weight/volume) for ½ MS. |
| Procedure Step 15 | Seeds are contaminated | Seeds were not sterilized properly | Use fresh seed sterilization solution. |
| | | | Use fresh 80% ethanol<br>Reduce the number of seeds per tube to ensure adequate exposure to the washing solution. |
| Part 4 Step 15 | Poor seed germination | Seed sterilization was too harsh. | Reduce incubation time in the seed sterilization solution. Make sure the colours of the seeds are not bleached. |
| | | | Immediately rinse off ethanol |
| | | Residual seed sterilization solution or ethanol remained on the seeds. | Repeat the rinsing step (Box 1 Step 10) |
| Part 4 Step 15 | Microbial colonies not observed on plates | Soil microbes no longer viable, ½ MS medium was too hot. | Allow the ½ MS medium to cool to 40°C before adding soil inoculum (Part 3 Step 7). |
| | | pH of the ½ MS medium was not suitable for required bacterial communities. | Adjust the pH of the ½ MS medium to match the pH conditions of the corresponding soil sample or test a range of pH values. |
| Part 4 Step 15 | Low diversity of visible colonies | Insufficient amounts of 10% AS were prepared. | Prepare at least 50 mL of 10% AS to avoid bias towards only high-abundance microbes. |

| | | | |
|---|---|---|---|
| Part 4 Step 15 | Contaminations on ArtSoil | 10% AS was not allowed to cool down between autoclaving steps. | Let 10% AS sit at RT for at least 16 h before the next autoclave. |
| Part 4 Step 15 | Contaminations on ArtSoil | ½ MS was contaminated. | Use freshly prepared ½ MS medium. Make sure ½ MS medium did not sit overnight prior to autoclave. |
| | | ArtSoil contains sugar. | Make sure to use MS powder that does not contain any sugars. |
| | | Contamination from working environment. | Make sure to prepare 50% FS, the ASE dilutions and ArtSoil in a biosafety cabinet. |
| | | Petri dishes were not sealed properly. | Seal the petri dishes properly before taking out of the biosafety cabinet. |
| | | | Use wider Micropore tape or seal the petri dishes twice. Perform pressure on the Micropore tape especially at the corners of the petri dish. |
| Part 4 Step 15 | ArtSoil dries out after prolonged incubation | Insufficient amount of ArtSoil media per petri dish. | Adjust the volume of ArtSoil according to the duration of your experiment. Increase the volume for experiments that require longer incubation periods or elevated temperatures in the growth chamber. |
| | | Petri dishes were not sealed properly. | Use wider Micropore tape or seal the petri dishes twice. Perform pressure on the micropore tape especially at the corners of the petri dish. |

<u>Timing</u>

Box 1: Seed preparation, 3-5 days
Box 2: Preparation of ½ MS, 1 day
Part 1: Preparation of 10% AS, 3 days
Part 2 and 3: Preparation of ASE and ArtSoil media, 30 mins
Part 4: Seed sowing, 30 mins (depending on quantity of seeds)
Box 3: Preparation of open system ArtSoil, 1 hr.
